## Appendix Tables S1 and S2, Appendix Figure S1 for "Recognition of phylogenetically diverse pathogens through enzymatically amplified recruitment of RNF213"

**Table of content**

| | page |
|---|---|
| Appendix Table S1 | 2 |
| Appendix Table S2 | 4 |
| Appendix Figure S1 | 6 |

### Appendix Table S1

Maximum likelihood analysis of positive selection among simian primate RNF213 sequences using the codeml algorithm. Shown are the results from a full-length alignment (top row), and from 6 alignments that each represent a segment of RNF213 that is free from recombination according to the GARD algorithm. For each alignment, we present the overall 'average' dN/dS (codeml model 0), the p-values for two complementary tests for positive selection (model 8 versus 8a, and model 8 versus 7), and predictions from model 8 of the proportion and dN/dS of rapidly evolving sites. Our findings are robust to the use of different starting parameters: we show results for four parameter combinations: initial_omega=0.4 or 3, and codon_model=2 or 3.

| Alignment | Codon model | Initial omega | Segment start nucleotide position in alignment | Segment end nucleotide position in alignment | Alignment length (codons) | Number of sequences | Overall dN/dS (model 0) | p-value model 8 versus 8a | p-value model 8 versus 7 | Percent of sites under positive selection | Estimated dN/dS of sites under positive selection | Number of sites under positive selection (BEB posterior probability >=0.9) |
|---|---|---|---|---|---|---|---|---|---|---|---|---|
| **Parameter set 1: codon model 2, initial omega 0.4** | | | | | | | | | | | | |
| full length | 2 | 0 | | | 5241 | 24 | 0,41 | 8,1E-42 | 8,0E-50 | 4,2 | 3,3 | 59 |
| GARD segment 1 | 2 | 0 | 1 | 1152 | 384 | 22 | 0,86 | 3,8E-05 | 7,4E-05 | 17,1 | 2,6 | 3 |
| GARD segment 2 | 2 | 0 | 1156 | 1827 | 224 | 23 | 0,85 | 3,5E-16 | 3,6E-16 | 7,6 | 6,3 | 11 |
| GARD segment 3 | 2 | 0 | 1831 | 4260 | 810 | 24 | 0,79 | 2,9E-16 | 2,8E-15 | 11,4 | 3,3 | 42 |
| GARD segment 4 | 2 | 0 | 4264 | 10791 | 2176 | 24 | 0,24 | 1,1E-09 | 2,9E-12 | 2,9 | 2,8 | 18 |
| GARD segment 5 | 2 | 0 | 10795 | 14217 | 1141 | 24 | 0,44 | 3,8E-10 | 3,9E-10 | 3,7 | 3,7 | 13 |
| GARD segment 6 | 2 | 0 | 14221 | 15723 | 501 | 24 | 0,30 | 1 | 1 | | | |
| **Parameter set 2: codon model 2, initial omega 3** | | | | | | | | | | | | |
| full length | 2 | 3 | | | 5241 | 24 | 0,41 | 8,1E-42 | 8,0E-50 | 4,2 | 3,3 | 59 |
| GARD segment 1 | 2 | 3 | 1 | 1152 | 384 | 22 | 0,86 | 3,8E-05 | 7,4E-05 | 17,1 | 2,6 | 3 |
| GARD segment 2 | 2 | 3 | 1156 | 1827 | 224 | 23 | 0,85 | 3,5E-16 | 3,6E-16 | 7,6 | 6,3 | 11 |
| GARD segment 3 | 2 | 3 | 1831 | 4260 | 810 | 24 | 0,79 | 2,9E-16 | 2,8E-15 | 11,4 | 3,3 | 42 |
| GARD segment 4 | 2 | 3 | 4264 | 10791 | 2176 | 24 | 0,24 | 1,1E-09 | 2,9E-12 | 2,9 | 2,8 | 18 |
| GARD segment 5 | 2 | 3 | 10795 | 14217 | 1141 | 24 | 0,44 | 3,8E-10 | 3,9E-10 | 3,7 | 3,7 | 13 |
| GARD segment 6 | 2 | 3 | 14221 | 15723 | 501 | 24 | 0,30 | 1 | 1 | | | |

### Appendix Table S1 (continued)

| Alignment | Codon model | Initial omega | Segment start nucleotide position in alignment | Segment end nucleotide position in alignment | Alignment length (codons) | Number of sequences | Overall dN/dS (model 0) | p-value model 8 versus 8a | p-value model 8 versus 7 | Percent of sites under positive selection | Estimated dN/dS of sites under positive selection | Number of sites under positive selection (BEB posterior probability >=0.9) |
|---|---|---|---|---|---|---|---|---|---|---|---|---|
| **Parameter set 3: codon model 3, initial omega 0.4** | | | | | | | | | | | | |
| full length | 3 | 0 | | | 5241 | 24 | 0,42 | 2,4E-44 | 2,7E-52 | 4,5 | 3,3 | 59 |
| GARD segment 1 | 3 | 0 | 1 | 1152 | 384 | 22 | 0,92 | 5,0E-07 | 1,7E-06 | 16,2 | 3,1 | 6 |
| GARD segment 2 | 3 | 0 | 1156 | 1827 | 224 | 23 | 0,85 | 6,0E-17 | 5,3E-17 | 7,3 | 6,7 | 10 |
| GARD segment 3 | 3 | 0 | 1831 | 4260 | 810 | 24 | 0,87 | 1,6E-19 | 1,6E-18 | 11,2 | 3,8 | 34 |
| GARD segment 4 | 3 | 0 | 4264 | 10791 | 2176 | 24 | 0,24 | 1,5E-09 | 1,8E-12 | 3,4 | 2,5 | 17 |
| GARD segment 5 | 3 | 0 | 10795 | 14217 | 1141 | 24 | 0,45 | 6,4E-11 | 9,1E-11 | 4,3 | 3,6 | 13 |
| GARD segment 6 | 3 | 0 | 14221 | 15723 | 501 | 24 | 0,31 | 1 | 1 | | | |
| **Parameter set 4: codon model 3, initial omega 3** | | | | | | | | | | | | |
| full length | 3 | 3 | | | 5241 | 24 | 0,42 | 2,4E-44 | 2,7E-52 | 4,5 | 3,3 | 59 |
| GARD segment 1 | 3 | 3 | 1 | 1152 | 384 | 22 | 0,92 | 5,0E-07 | 1,7E-06 | 16,2 | 3,1 | 6 |
| GARD segment 2 | 3 | 3 | 1156 | 1827 | 224 | 23 | 0,85 | 6,0E-17 | 5,3E-17 | 7,3 | 6,7 | 10 |
| GARD segment 3 | 3 | 3 | 1831 | 4260 | 810 | 24 | 0,87 | 1,6E-19 | 1,6E-18 | 11,2 | 3,8 | 34 |
| GARD segment 4 | 3 | 3 | 4264 | 10791 | 2176 | 24 | 0,24 | 1,5E-09 | 1,8E-12 | 3,4 | 2,5 | 17 |
| GARD segment 5 | 3 | 3 | 10795 | 14217 | 1141 | 24 | 0,45 | 6,4E-11 | 9,1E-11 | 4,3 | 3,6 | 13 |
| GARD segment 6 | 3 | 3 | 14221 | 15723 | 501 | 24 | 0,31 | 1 | 1 | | | |

**Appendix Table S2**

**CryoEM data collection and processing, model refinement and validation statistics.**

| | RNF213 (EMDB 19653) (PDB 8S24) |
|---|---|
| **Data collection and processing** | |
| Microscope | Titan Krios |
| Specimen temperature (K) | ~80 |
| Voltage (kV) | 300 |
| Camera | Falcon 4i |
| Energy filter | Selectris X |
| Energy filter slit width (eV) | 10 |
| Electron fluence (e–/Å$^2$) | 29.8 |
| Electron flux ($e^-$/pix/s) | 5.2 |
| Exposure time (s) | 4.86 |
| Magnification | 130,000× |
| Pixel size (Å) | 0.921 |
| Defocus range (μm) | 0.5 – 3.0 |
| Average defocus (μm) | 2.0 |
| Total number of movies | 7,022 |
| **Data processing** | |
| Initial number of particle images | 1,010,981 |
| Final number of particle images | 143,490 |
| Particle box size (pixels) | 512×512 |
| Symmetry imposed | C1 |
| Map resolution (Å) | 3.0 Å (0.143 global FSC cutoff) |
| Local resolution range (Å) | 2.5 - 4.5 Å |
| **Refinement** | |
| Initial model used (PDB code) | N/A |
| Refinement package | COOT, Phenix |
| Model resolution (Å) | 3.0 Å (0.5 map-model FSC cutoff) |
| Map sharpening *B* factor (Å$^2$) | 30 Å$^2$ |

| | |
|---|---|
| Model composition | |
| Non-hydrogen atoms | 35,357 |
| Protein residues | 4387 |
| Ligands | 3 (Mg, Zn, ATP) |
| Molecular weight (kDa, incl H) | 503 |
| *B* factors (Å$^2$) | |
| Protein | 90 |
| Ligand | 60 |
| RMS deviations | |
| Bond lengths (Å) | 0.01 |
| Bond angles (°) | 0.5 |
| Validation | |
| MolProbity score | 1.7 |
| Clashscore | 8.9 |
| Poor rotamers (%) | 0.7 |
| Cβ deviations (%) | 0.0 |
| CaBLAM outliers (%) | 1.7 |
| Ramachandran plot | |
| Favored (%) | 96.6 |
| Allowed (%) | 3.2 |
| Outliers (%) | 0.2 |

### Appendix Figure S1

Confocal micrographs of RNF213$^{KO}$ MEFs stably expressing the indicated RNF213 variants. Cells were fixed at 3.5h post-infection with mCherry-expressing *S.* Typhimurium, 6h post-infection with mCherry-expressing *L. monocytogenes* ΔActA and 24h post-infection with Tomato-expressing *T. gondii* Type I strain RH. Scale bar, 10µm.

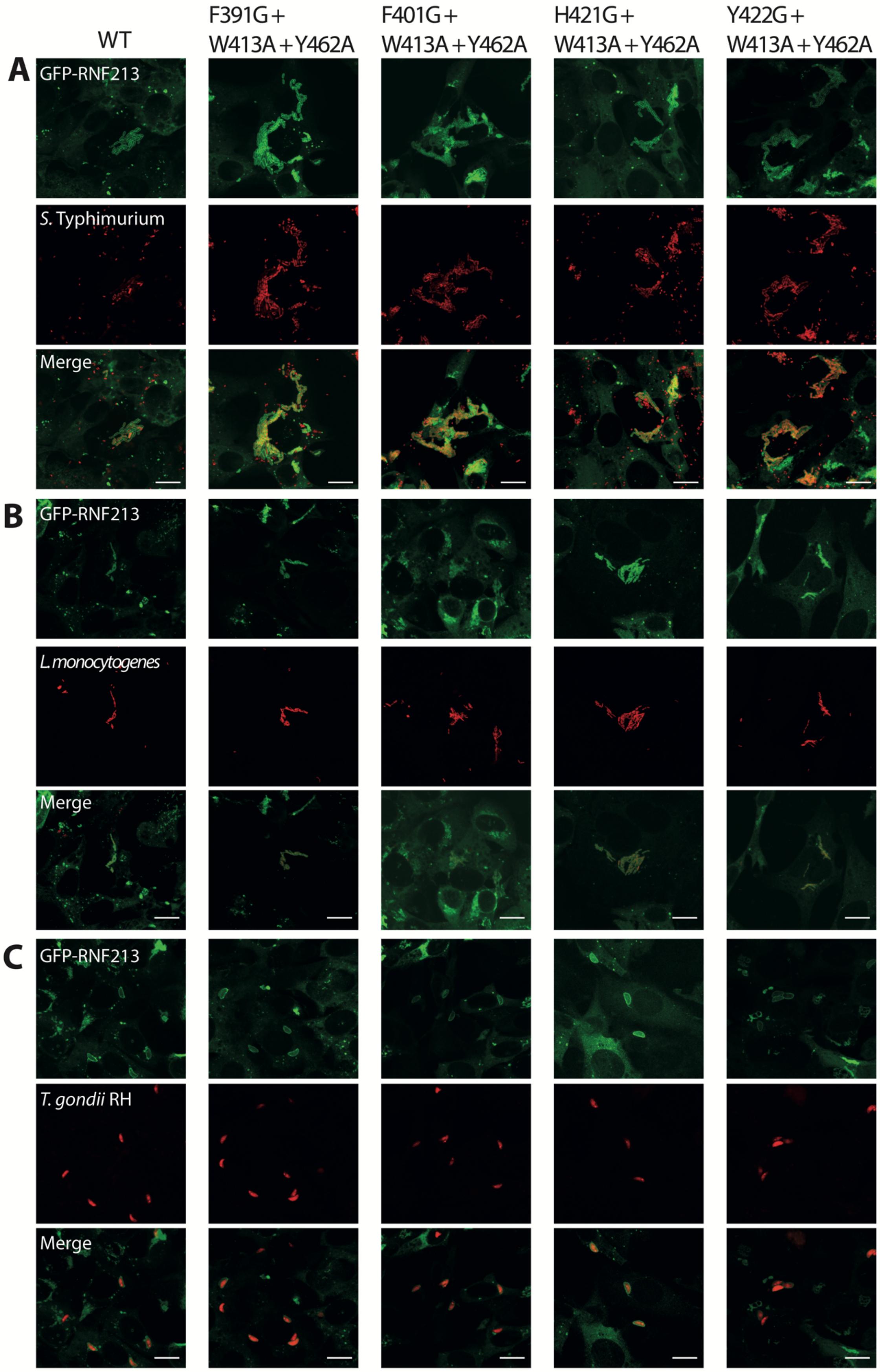
