## Supplementary material for "Recognition of phylogenetically diverse pathogens through enzymatically amplified recruitment of RNF213": Dataset EV2: readme_Dataset2.txt

Dataset 2.
Site-by-site results of positive selection analysis. For each codon in the alignment of simian primate RNF213 sequences, we present sitewise results from codeml�s model 8 analysis of positive selection, using either the full-length alignment (columns 5, 6 and 7) or shorter segments of the alignment that are free from recombination (columns 8-11), in each case presenting the Bayes Empirical Bayes posterior probability that each site is among the rapidly evolving subset, as well as codeml�s posterior mean estimate of dN/dS.
